# Leaving Academia: Insights from Evolutionary Biologists on Their Career Transitions and Job Satisfaction

**DOI:** 10.1101/2025.08.19.671061

**Authors:** Andrea J. Betancourt, Seth Barribeau, Hildegard Uecker, Svenja Hammer, Claire Asher

## Abstract

Many who have obtained PhDs in evolutionary biology will ultimately pursue careers that fall outside a narrow definition of an academic career. At the same time, PhD students and supervisors of PhD students are often ill-informed about career options outside of academia. Here, we report on a survey of evolutionary biologists who have pursued non-academic careers, to understand what careers they pursue, how they transitioned into those careers, how well prepared they were, and how satisfied they are with their current work. Overall, the message from this survey is positive– evolutionary biologists are readily employable outside of academia, generally well-prepared for those jobs, and report high levels of satisfaction in their non-academic careers. We also highlight areas where preparation for non-academic careers could be improved, which might be addressed by individual mentors or PhD training programmes.

## Introduction

Many PhD holders end up leaving the academic career track (Lu et al., 2023; National Center for Science and Engineering Statistics, 2023) – with the academic career track narrowly defined here as PhD graduate to postdoctoral researcher to independent principal investigator. One major reason is that while the rate of PhD training has increased over recent decades, the growth in academic positions has not kept pace (Organisation for Economic Co-operation and Development, 2021, 2023; Sarrico, 2022). Intentionally or not, then, the purpose of PhDs has expanded, from solely training academics to also training workers for other careers (Hancock, 2023; McAlpine, 2020, 2024; Sarrico, 2022). At the same time, academics are often poorly equipped to advise trainees on non-academic careers (McAlpine, 2024; Woolston, 2022a).

Against this backdrop, this study– a survey of those trained in evolutionary biology who now have non-academic careers– aims to provide a resource for PhD students and graduates exploring their career options, as well as for their academic advisors. Complementing large and general survey studies (e.g., Nordling, 2025), we specifically focus on biologists trained in evolutionary biology. Researchers with very different training and skill sets fall under this umbrella, so that training in evolutionary biology can be diverse, and suited to widely different career paths. Accordingly, young evolutionary biologist may be especially unaware of their nonacademic career options, as, for example, the alumni of their PhD program in evolutionary biology may not have pursued career paths relevant to them. A survey specifically among -- and for -- evolutionary biologists therefore may be especially useful.

Overall, with the caveat that any survey relying on voluntary responses will be subject to biases, we find an optimistic picture for evolutionary biologists seeking non-academic jobs. Our respondents found jobs quickly, in a diverse range of industries, and report high levels of satisfaction with their jobs. In addition, they use many of the skills they learned during their training– unsurprisingly, data analysis skills were particularly well-cited, but so were generalist skills that most trainees obtain regardless of their specialist area of training.

## Survey design

We surveyed evolutionary biologists about their experiences leaving the academic career track. The survey, which was delivered on the JISC online surveys platform (https://app.onlinesurveys.jisc.ac.uk/), consisted of 80 free text and multiple-choice questions, including 22 questions about non-academic jobs, 24 about academic training, 13 on the transition to the non-academic track, and 11 questions on satisfaction in and outside of academia (see Supplementary File). A further 10 questions concerned personal background information of the participants, including gender and location. Anonymous responses were solicited via advertising the survey on social media, through broadcast emails (*e*.*g*., via professional societies and the evoldir mailing list), and through direct contact.

## Surveyed population

We obtained 178 total responses. Eight responses were eliminated due to being mostly incomplete (*n* = 2), consent to use the data was not given (*n* = 4), or respondents had not started a PhD (*n* = 2, too few to be representative of this cohort). We analyzed the remaining 170 responses (summarised in Supplementary Tables), mainly descriptively.

Our survey respondents identified as 54.1% female, 44.1% male and 1.2% other (out of 169 responses). At the time of the survey, they were working in 23 countries, with roughly half in either the United States (29.4%) or the United Kingdom (20.6%), and almost all in either North America or Europe. They appeared to be globally mobile, with 37.8% not living in their country of origin at the time of the survey, and just over half (53.0%) having lived in at least two countries. Many may have moved to pursue training, as their country of origin differs from the country where they obtained their PhD (37.4%), or moved during training (e.g., between PhD and postdoc, 46.2%), consistent with the mobility of STEM PhDs students overall (Woolston, 2022b).

Most of the respondents had left the academic career track in the recent past, within the last 5 years (59.4%), and left either immediately after their PhD (24.9%), or during the early postdoc years (≤ 5 years; 39.6%; **Figure 1a and b**). Because there is concern that the pipeline to permanent jobs is especially narrow for female academics, we analysed these data by gender, finding a non-significant trend for women to leave earlier than men (χ^2^ = 9.91, df = 5, *p* = 0.078). This result is broadly consistent with a large, multidisciplinary study which found that, in biology, women stop publishing scientific papers earlier than men on average, though the gender difference had diminished in recent years (Kwiek & Szymula, 2025). Respondents were trained in a broad range of subfields within evolutionary biology, with the two most common subfields being ‘Evolutionary Ecology’ and ‘Population and Quantitative Genetics’ (**Figure 1c**).

**Figure 1.**
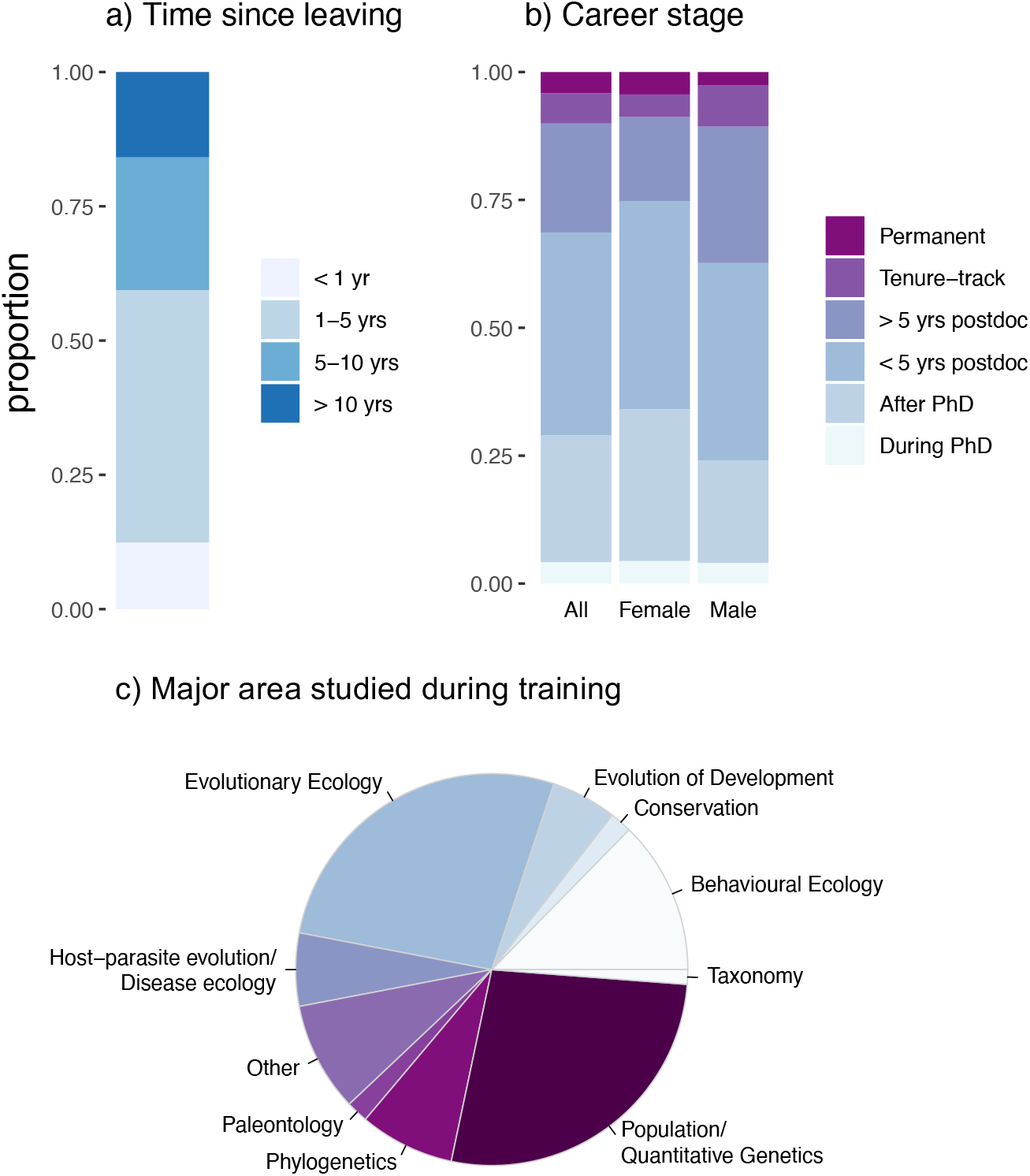
Characteristics of the surveyed sample. **a)** The length of time prior to the survey and **b)** career stage at which respondents for this survey left the academic career track **c)** The subfield of evolutionary biology studied during training.

## Limitations

Several caveats and limitations apply to these results. Our analysis is limited to a particular cohort of individuals—people who left the academic career track at some point after starting a PhD in evolutionary biology or a closely related field. Survey responses were voluntary and thus may well have come disproportionately from those who felt positively about their non-academic careers. Due to how this survey was disseminated—largely through professional academic networks-- our results about when people left academia may not be representative. In particular, our survey may have been unlikely to reach those who left very early (e.g., before obtaining a PhD), or those who left the academic track long ago. However, the experiences of those who have recently left academia are probably most relevant to those considering a move into a non-academic career. And, necessarily, our results are retrospective, with the possibility of rapid changes to the career landscape driven by, e.g., generative artificial intelligence, and recent upheavals affecting U.S. science funding.

## Survey Results

### Leaving academia– why

Respondents had a choice of up to seven categories of reasons for leaving academia (**Table 1**). Most people did, in fact, cite more than one category (87%), with a median of three reasons chosen, suggesting that the reasons for leaving academia are multifaceted. That said, the top two reasons were ‘lack of job opportunities’ (62.9%) and ‘lack of job security’ (60.0%), and had significant overlap (*n* = 75; Fisher’s exact test, *p* = 0.00063). Together with the result that people tend to leave at career stages when employment is precarious, these results suggest that the primary reason most people leave academia is due to the scarcity of permanent academic job opportunities. To put these results in context, it is important to note that job security can also be a concern outside of academia, with jobs sometimes lost at shorter notice than is usual in academia, and not many jobs with the permanence equivalent to tenured academic jobs. However, maintaining an academic career usually requires securing an unbroken series of academic jobs, often accompanied by a move, until a rare permanent academic position is acquired. This entanglement of job insecurity with a shortage of positions and a need to relocate is likely what drives many researchers to leave the academic career track.

**Table 1.** Reasons cited for leaving academia, out of 544 total reasons from 170 respondents.

| Reason | count |
| --- | --- |
| Lack of job opportunities within academia | 107 |
| Lack of academic job security | 102 |
| Geographic constraints/didn't want to or couldn't move | 70 |
| Preferred non-academic work environment or non-academic work | 65 |
| Financial reasons | 61 |
| Personal/Family reasons | 58 |
| Physical or mental health reasons | 56 |
| Other | 25 |

Nevertheless, most individuals who cited job opportunity/security also cited other reasons (*n* = 59 of 75), suggesting that while job opportunities are important, they are not the only reason people leave. The remaining reasons for leaving academia were cited more or less evenly, by between 30-40% of respondents each. The only positive reason given as an option– a preference for non-academic work– was the fourth most highly cited, but was usually cited in conjunction with other reasons (n = 58 of 65 cases), suggesting that most were pushed, rather than pulled, out of academia. The reasons cited by gender differed for men and women (χ^2^ = 28.99, df = 7, *p* = 0.00015), mainly as women more often cited “mental or physical health” as a reason for leaving (*n* = 39/294 vs. 16/239, Pearson’s adjusted residual = 4.35, *p* = 6.547e-06; Bonferroni-corrected α = 0.00625).

### Leaving academia– how

We asked several multiple-choice questions about the search for their first post-academic job. We asked how respondents found their first academic job (**Figure 2A**). The strategies used to find jobs were diverse. In keeping with previous studies (e.g.,Germain-Alamartine et al., 2021) networking was found to be useful in transitioning to a non-academic job. A substantial fraction made use of non-academic connections (23.1%), and somewhat contrary to our expectations, an even larger fraction leveraged some sort of academic connection (34%). Many of the remaining strategies were conventional job search strategies, such as applying through websites. A few were recruited or started their own businesses. For most respondents, the job search appears to have been straightforward. Most prepared their documents on their own without assistance (73.3%). The majority (77.1%) obtained a job within six months, nearly half within three, with fewer than 10 applications (72.1%) and without additional training (75.8%; **Figure 2B-D**).

**Figure 2.**
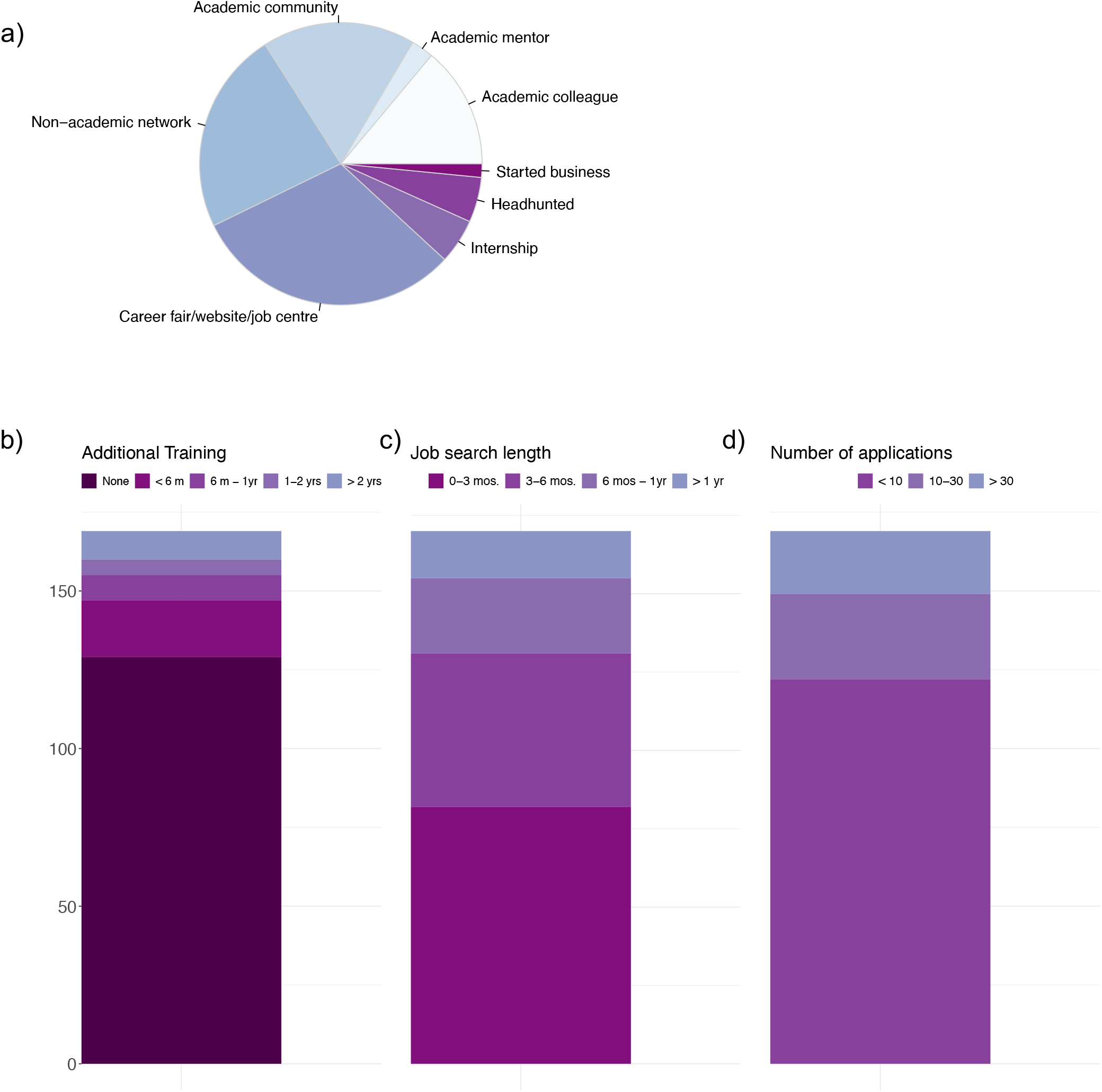
The survey asked how respondents about their first non-academic job search. Responses are multiple choice with the prompt: a) How did you find your first job outside of academia?, b) How many jobs did you apply for before you found your first job? c) How long in advance did you start looking for non-academic jobs before you found your first job? and d) How many jobs did you apply for before you found your first job? Answers to a) had a free text ‘other’ category; the free text responses were manually curated and are included here (n = 194); others were single answer only. One answer to b) had two responses; we used the longest time period given.

### Non-academic careers for evolutionary biologists

We asked about the types of jobs currently held by people trained in evolutionary biology. We reasoned that many respondents would be likely to be working either in education or in scientific research outside of our narrow definition of the standard academic career track, and therefore asked additional questions of these respondents. Somewhat contrary to this expectation, a relatively modest number of the respondents work in education (24 of 166); we did not further break down these responses due to their limited number. Many of the respondents, however, (55.9%) were working in scientific research, at least partly.

For those working in scientific research, we broke down the responses in several ways (**Figure 3**). We asked about the work they did. Many indicated they worked doing some form of data analysis and/or empirical research (*n* = 70 and 49, respectively). We asked about the organisations they worked for; most (52.6%) work at least partly for for-profit businesses, with the remainder split rather evenly over other kinds of organizations. We asked them to describe the setting of their work; ‘dry laboratory’ or ‘office (management/administration)’ were the most popular choices (*n* = 59 and 58, respectively), but a similar number were engaged in work in the field or laboratory (*n* = 52). Finally, we asked for 3-5 keywords describing their current work. While difficult to fully summarize the responses, as there were over 400 unique keywords, many of the respondents appeared to be employed in some form of data analysis. Among the most frequent responses were keywords similar to bioinformatics (*n* = 17), genomics (*n* = 16), or some other form of data analysis (data: *n* = 47, statistics: 9 [not also including ‘data’], or machine learning, *n* = 8). Similarly, there were responses that indicated careers using computational or programming skills (software and computation, *n* = 8 each).

**Figure 3.**
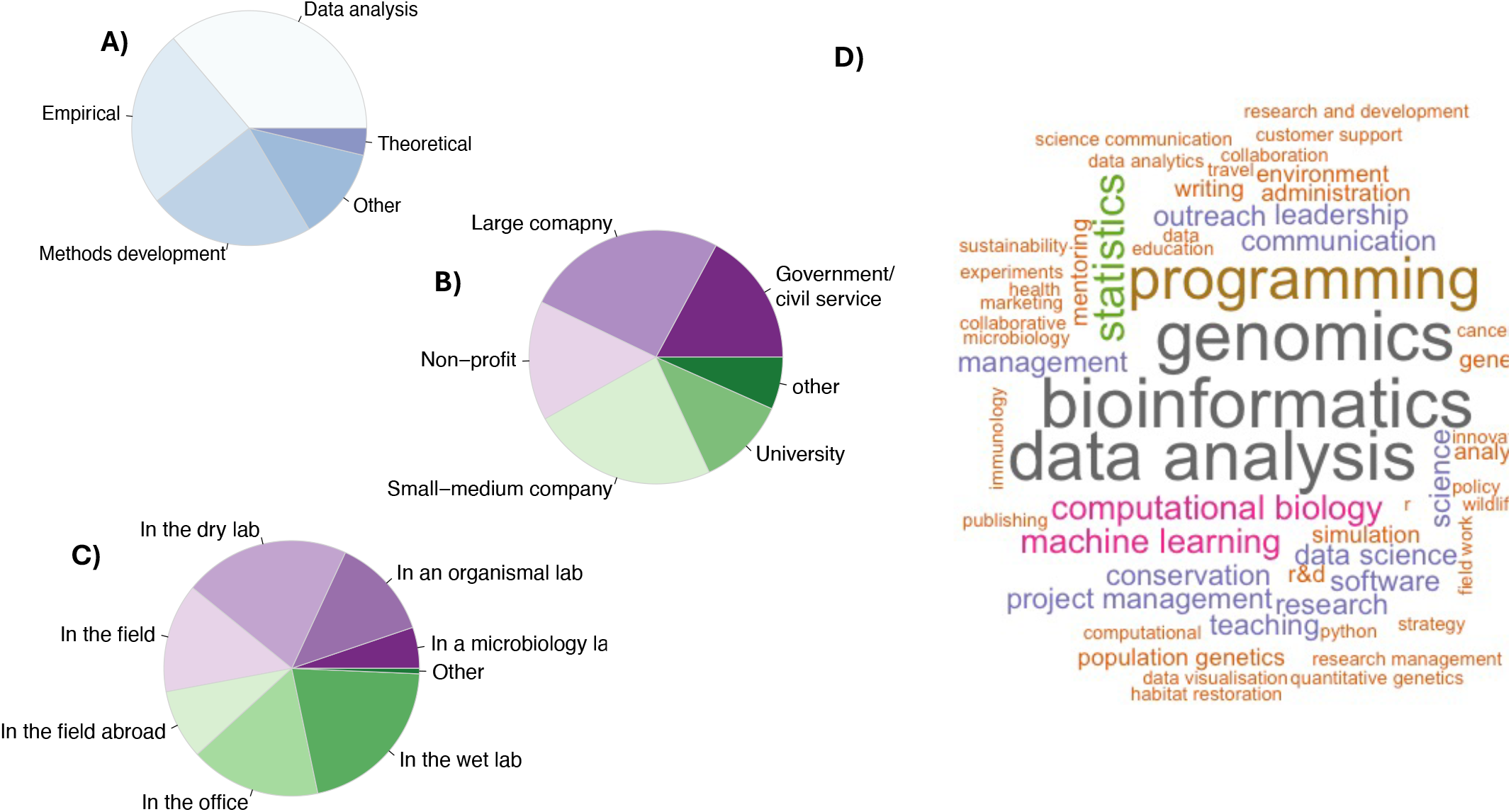
Summaries of answers to several questions about non-academic jobs in scientific research, with respondents asked about aspects of their current job, including A) the type of work B) the type of organization that they work for C) the setting of their work, and D) a description of their current job. A-C were multiple choice questions; D was free text.

For those working in neither education nor research, we obtained a further 80 free text responses, and roughly categorized these into industry manually. Consulting and the tech industry comprised the largest number of these (*n* = 13 each), closely followed by jobs in the public sector in some form (*n* = 10), writing/publishing (*n* = 7) and the biotech industry (*n* = 6).

### Are evolutionary biologists prepared for non-academic careers

We were interested in knowing how well-prepared evolutionary biologists were for their current jobs, based on their skills used in their jobs and those obtained during the PhD/postdoctoral training. We asked about 22 types of skills, falling into data collection/lab skills, analysis, and general (different types of writing, researching literature, etc.) categories. Overall, it appears that evolutionary biologists are well-prepared for their current jobs – individuals reported that they were trained in skills necessary for their job more often than not (**Figure 4**, χ^2^ =220.02, df = 12, *p* < 2.2e-16), particularly in statistics and scientific writing (adjusted Pearson’s residuals = 4.47 and 5.05, *p* < 0.05). Consistent with these matches between skills and jobs, most people self-reported that they felt prepared for their jobs (**Figure 5A**, median = 3, with 1 = not at all and 4 = well-prepared). This feeling of preparation did not differ based on the stage at which training ended (*r*_*s*_ = 0.111, *p* = 0.153), or the gender of the respondent (Mann Whitney U test, *p* = 0.493) in this data set. Further, the number of skills cited did not appear to depend on career stage achieved (*r*_*s*_ = 0.880, *p* = 0.253).

**Figure 4.**
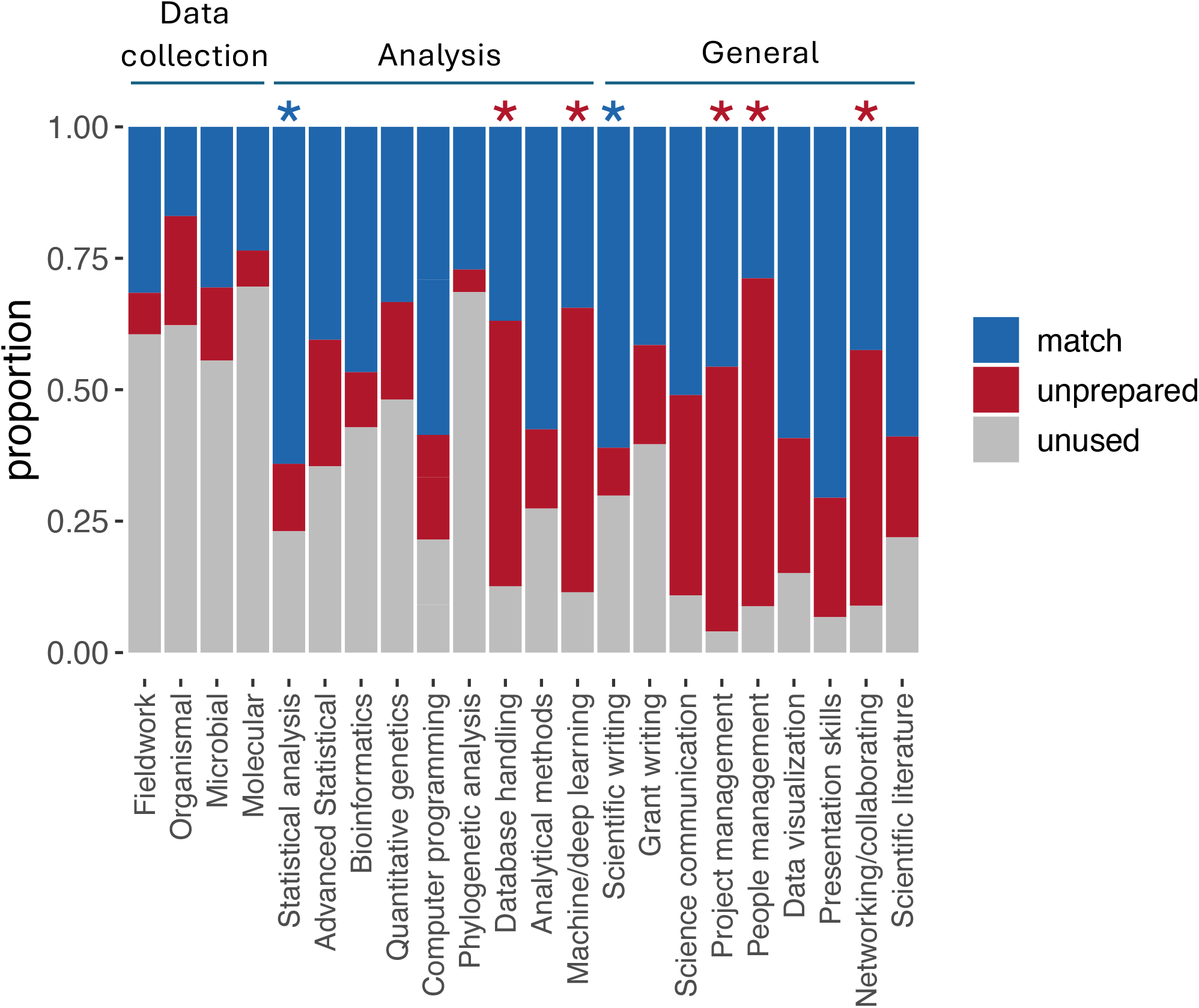
Match in skills between training and jobs. These are based on the answers to a series of multiple-choice questions asking about skills acquired during PhD and postdoctoral training, either informally or formally, and used in respondents’ current jobs. Bars are normalized to show only respondents that selected a skill in either training or jobs, and show the proportion of these that selected the skill for both training and job (‘match’), for training only (‘unused’), and for the job only (‘unprepared’). Overall, there were more cases where a skills used on the job and acquired in trainig were matched (see text); stars show categories where adjusted Pearson’s residuals indicated significant differences between after Bonferroni correction (with blue indicating more matches than mismatches, and red fewer).

**Figure 5.**
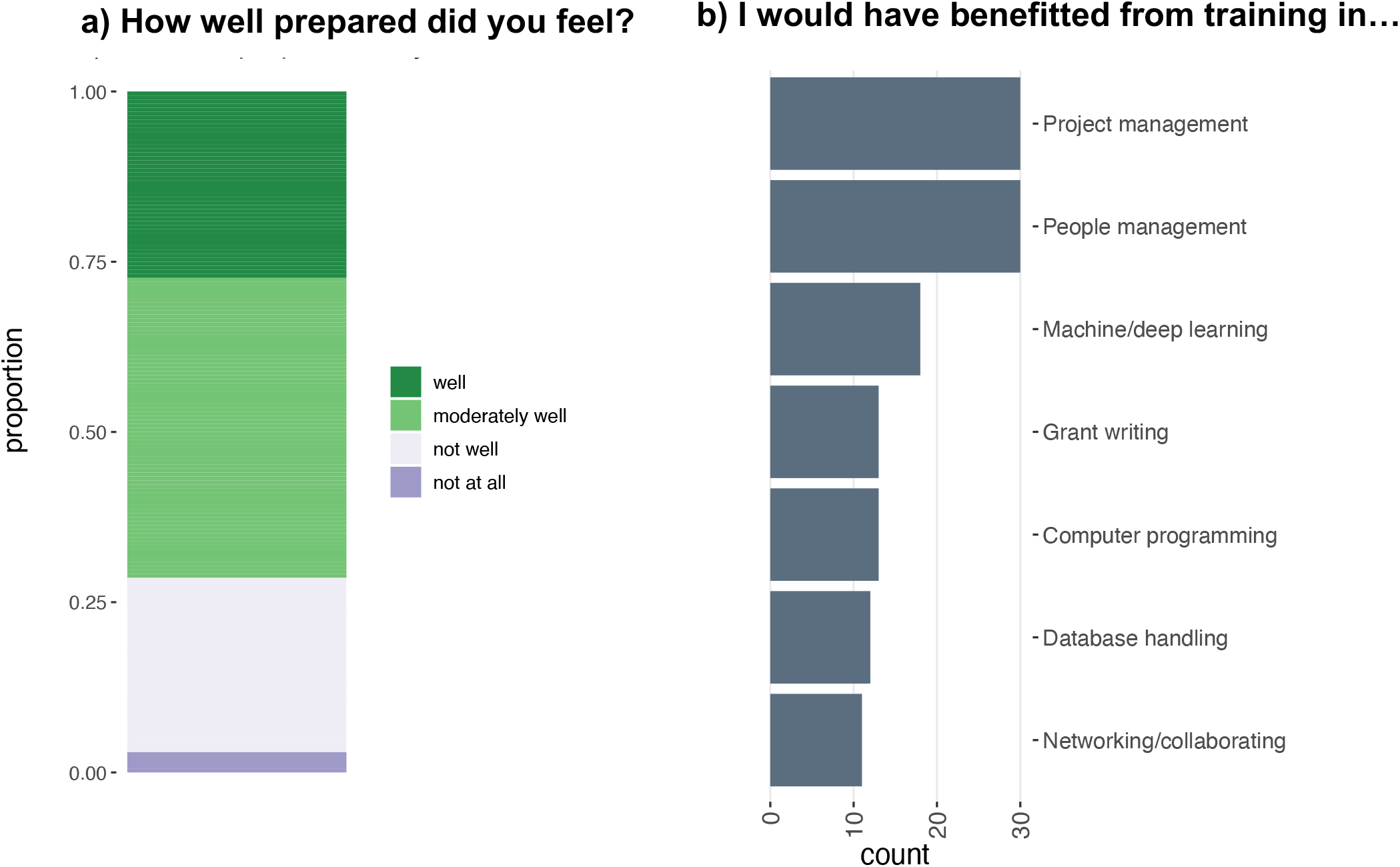
Preparation for non-academic careers. Respondents were asked A) to rank how well they felt their training prepared them for their first non-academic job on a four-point scale B) what skills they wish they had acquired during their training. Shown are those indicated in 10 or more responses.

There were, however, several areas where there were mismatches between training and skills required on the job. Data collection skills, in particular, were mostly unused. That said, the underlying knowledge that comes in training in data collection skills might be used indirectly, such as in cases where the job involves management of data production teams, or analysis of the data produced. Unfortunately, we did not ask specifically about this kind of application of skills. Importantly, there were some training gaps, especially in ‘machine learning’ and ‘database handling’ among the analysis skills, and project and people management under general skills (**Figure 5A)**. Unsurprisingly, these skills were among the top skills where people expressed a desire for more training when prompted (**Figure 5B**). That people felt they would have benefitted from project management training was surprising, as academic research is largely managing projects. It may be that respondents refer specifically to formal certification in project management, or it may simply be that managing projects is a complex task that people feel they could be better prepared for.

### Career satisfaction inside and outside academia

We asked questions pertaining to different aspects of satisfaction. These included three pairs of questions designed to compare satisfaction inside *vs*. outside of academia *(i)* overall, *(ii)* financially, and *(iii)* at attaining work-life balance. For all three categories, the respondents reported significantly higher job satisfaction in their current jobs than in their academic positions (**Figure 6A**; Mann-Whitney U test, *p* < 2.2 e -16 for all three comparisons). Interestingly, respondents that felt better prepared for their jobs also reported higher overall job satisfaction (*r*_*s*_ = 0.180, *p* = 0.019).

**Figure 6.**
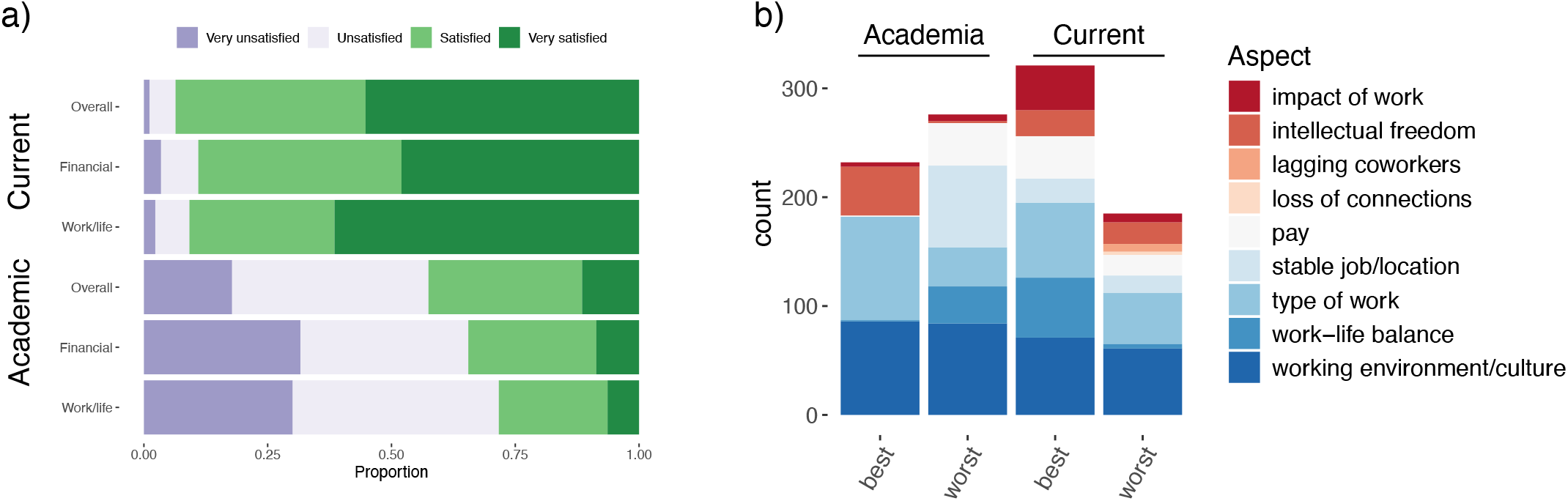
**a)** Satisfaction on a four category Likert scale for overall, financial, and work-life balance aspect of current (top) and academic (bottom) careers. **b)** Free text responses to questions about factors that most influenced satisfaction, positively (“best”) or negatively (“worst”) in academia and on the job, coded into categories. At least one of these questions was answered by 129 survey participants, which we coded into 1014 responses (*n* = 232, 276, 321, and 185 for the academia best and worst, and current best and worst, respectively).

To understand what factors drive satisfaction in these broad categories, we asked for free-text descriptions of factors that most contributed to satisfaction, positively or negatively, in academia and non-academic jobs, and manually coded these responses into categories to capture general patterns (**Figure 6B)**. Working environments were rather evenly, and highly, cited across both academic and non-academic jobs as contributing both favorably and unfavorably to satisfaction. That said, responses that suggested or mentioned abuse, harassment, and bullying, almost all came from within academia (*n* = 14 vs. *n* = 2). We have little additional context in most of these cases, but the problems in academia have been addressed elsewhere (Björklund et al., 2021; Jones et al., 2019; Manuel et al., 2023; Offer et al., 2023; Woolston, 2022b). Other strongly negative assessments of academic working environments cited stress, pressure, and long-working hours for little pay, consistent with large, multidisciplinary surveys of current STEM PhDs (Woolston, 2022c).

Within academia, respondents appeared to enjoy intellectual freedom (consistent with Woolston, 2019; *n* = 45), and the work itself (*n* = 69), but not job and location instability (*n* = 75). In their jobs, roughly similar numbers of respondents appeared to enjoy and dislike their work (*n* = 69 vs. 47). They also appreciated their work-life balance (*n =* 55) and having a direct impact with their work (*n* = 41). Perhaps tellingly, there were the fewest number of total responses recorded for the worst aspects of the job.

## Lessons learned

Overall, our results paint an optimistic picture for evolutionary biologists pursuing non-academic careers. On average, they found jobs quickly, without additional training, felt well-prepared for those jobs, and have high levels of overall satisfaction. This generally optimistic message is consistent with the fact that most students feel a PhD improves their job prospects (Woolston, 2022a). While some missed the intellectual freedom they had within academia, many appear to enjoy their non-academic work, particularly when it leads to rapid impact, as well as the stability, pay, and work-life balance that comes with a non-academic career. Despite the limitations of this survey, these results do show that training within academia can be valuable, and that leaving academia can be a positive experience.

Through providing mentors and trainees information about how academic training can contribute to a successful career outside of academia, we hope to contribute to a culture of frank discussion of non-academic career prospects (c.f., Goodwin et al., 2015; Organisation for Economic Co-operation and Development, 2023). While overall respondents felt well-prepared for a non-academic career, there were skills gaps for some types of data analysis, such as machine learning, and in some general areas, particularly people management and project management. Where there are gaps in formal training, PhD programmes or short courses, rather than individual supervisors, might be better placed to fill them. But mentors may be able to provide their mentees relevant experience— e.g., opportunities to manage people, present their work, write small grants— and communicate that this experience pays dividends for both academic and non-academic careers.

Finally, we would also like to point out the societal benefits of academic training: PhDs taking roles outside of academia may bring valuable skills and experience otherwise difficult to obtain (Santos et al., 2016). Our survey respondents took on protracted apprenticeships, presumably out of a keen interest in evolutionary biology, and, as a result, companies and governments can hire staff that have already achieved high levels of specialized expertise.

## Acknowledgments

We would like to thank the European Society of Evolutionary Biology (especially Nikki Cook and Ute Friedrich), the Society for Molecular Biology and Evolution, the American Society of Naturalists for promoting this survey. We would also like to thank all the respondents--every one of your answers was carefully read, even if we couldn’t represent them all fully due to privacy concerns. AJB was supported by ERC Consolidator Grant TE_INVASIONS.

## Notes

**Conflict of Interest** We have no conflict of interest to declare.

### Competing Interest Statement

The authors have declared no competing interest.

